# Stethoscope disinfection is rarely done in Ethiopia: what are the associated factors?

**DOI:** 10.1101/474098

**Authors:** Biniyam Sahiledengle

## Abstract

**Background:** The stethoscope, which is universally used as a medical device by healthcare providers, is likely to be contaminated by pathogenic microorganisms. And regular cleaning of the diaphragm of the stethoscope with a suitable disinfectant is decisive. However, in the resource constrained setting like many healthcare facilities in Ethiopia healthcare provider’s stethoscope disinfection practice and its associated factors have not been well studied so far. Therefore, this study sought to determine stethoscope disinfection practice and associated factors among the healthcare providers in Addis Ababa, Ethiopia.

**Methods:** A facility-based cross-sectional survey was carried out between April and May 2016. For this survey, 576 healthcare providers were included from 21 healthcare facilities in Addis Ababa. A pre-tested structured questionnaire was used for data collection. Descriptive statistics were computed. Binary and multivariable logistic regression analyses were used to identify factors that were significantly associated with stethoscope disinfection after every use.

**Results:** Five hundred forty six participants were take part in this study, for a response rate of 94.7%. Of these, only 39.7% (95%CI: 35.9, 44.0%) of healthcare providers disinfecting their stethoscope after every use. Physicians were less likely to disinfect there stethoscope compared to nurses (AOR=0.21; 95%CI: 0.09, 0.49). Healthcare providers who had awareness on infection prevention guideline, healthcare providers who had favorable attitude towards infection prevention and participants having safe infection prevention practice have better stethoscope disinfection practice after every use as compared to their counterparts (AOR=1.93; 95%CI: 1.31, 2.82), (AOR=1.73, 95%CI: 1.02, 2.93), and (AOR=3.79, 95%CI: 2.45-5.84), respectively.

**Conclusions:** Only a small proportion of healthcare providers disinfect their stethoscopes after every use. Factors such as awareness on infection prevention guidelines, favorable attitude towards infection prevention and safe infection prevention practice were the independent predictors of stethoscopes disinfection after every use. Hence, implementation of effective training on stethoscope disinfection along with increasing awareness on infection prevention may improve stethoscope disinfection practice.

## Background

Healthcare-associated infections (HAIs) continues to be one of the most important public health problems in many countries throughout the world [1,2], and remain a common global challenge mainly in low and middle-income countries [3]. Further, HAIs result in protracted hospital stays and increased substantial cost for healthcare system [4, 5]. It is known long ago HAIs are caused by viral, bacterial, and fungal pathogens. And many healthcare associated bacterial pathogens may well survive or persist on different surfaces for months and can thereby be a continuous source of transmission, if no regular preventive surface disinfection is performed [6,7]. Investigators of a systematic review also reported highly virulent microorganisms, particularly those known to cause nosocomial infections in admitted patients, such as *Enterococcus* species, *methicillin-resistant Staphylococcus aureus* (MRSA), *Escherichia coli*, *Klebsiella* species, *Pseudomonas aeruginosa*, and *Acinetobacter* species, are capable of surviving for several days on hospital surfaces [7]. Consequently, medical equipment surfaces, such as stethoscopes can easily contaminated with these infectious agents and contribute to the spread of HAIs [8,9,10,11,12], and may possibly cause outbreaks of hospital-acquired infections [13].

In this regard, the stethoscope, which is universally used as a medical device by healthcare workers, is more likely to be contaminated by microorganisms, if it is not disinfected and may transmit pathogens from one patient to another [14-18]. Without proper disinfection, stethoscopes represent a potential means of transmission of microorganisms from one patient to another, and between animate and inanimate objects and vice versa [18,19]. For this reason, regular cleaning with a suitable disinfectant is decisive and part of a multi-barrier strategy to prevent HAIs; since failure to properly disinfect or sterilize equipment carries not only risk associated with breach of host barriers but also risk for person-to-person transmission and transmission of environmental pathogens (e.g., *Pseudomonas aeruginosa*) [14, 20].

There is strong scientific evidences documenting the various pathogens found colonizing stethoscope surfaces [14,15,21,22]. Several studies have demonstrated that stethoscope membranes harbor bacteria, including resistant microorganisms such as Methicillin-resistant *Staphylococcus aureus* (MRSA), vancomycin-resistant *Enterococci* (VRE) and *Clostridium difficile* [14,23,24]. It was also reported that *S aureus* colonies can survive on stethoscope membranes for longer than 18 hours [25]. Moreover, highly resistant bacteria, MRSA can potentially survive up to 9 days on stethoscopes [26].

Effective disinfection of stethoscopes with isopropyl alcohol eliminates up to 99% of bacteria [22]. However, in the resource constrained setting like many healthcare facilities in Ethiopia, healthcare workers infection prevention compliance [27,28,29], and instrument disinfection practice was poor [30]. Consequently, the prevalence of HAIs was high [31,32,33]. There is also significant bacterial contaminations of stethoscopes were reported [34,35,36]. For example, a study by Shiferaw et al [34] reported that, of the 176 stethoscope examined 85.8% were contaminated and of 256 bacteria isolates, 52% were potential pathogens like *S.aureus, Klebsiella spp., Citrobacter spp., P.aeruginosa* and *E.coli*. Although, this evidence is available, there has still been a lack of studies about stethoscope disinfection in Ethiopia to overcome the problem. All the previously conducted related studies are case studies in one or similar type of hospital [34,35,36]. As well as none of the previously conducted studied include health centers in their assessment, where the majority of population seek out healthcare service. To the best of my knowledge this study is the most extensive study, investigating stethoscope disinfection practice in both hospitals and health centers. In order to improve healthcare provider’s appropriate disinfection practice and to develop successful infection preventive programs in-depth understanding of the issues is essential. Therefore, this study aimed to assess the practice of stethoscope disinfection among healthcare providers working in healthcare facilities in Addis Ababa, Ethiopia. Furthermore, the study aimed to identify factors associated with stethoscope disinfection. The findings of the study will used as an input for policy makers, programmers and healthcare workers to improve quality services and to prioritize interventions by decision makers to overcome the problem.

## Materials and Method

### Study area and design

A facility based cross-sectional study was conducted from April 11 to May 20, 2016 in randomly selected 21 healthcare facilities found in Addis Ababa (the capital city of Ethiopia). Administratively, the city is divided in to 10 sub-cities and 116 districts. Addis Ababa's population was estimated to be 3,273,000 in 2014/15, of which 1,551,000 (47.4%) were males and 1,722,000 (52.6%) were females [37]. At the time of this study there were a total of 90 health centers and 13 public hospitals were found in Addis Ababa. In these healthcare facilities around 7,642 healthcare providers were working at the time of this study.

### Study participants

All healthcare providers found in public hospitals and health centers in Addis Ababa were the source population. Selected healthcare providers who work at least 6 months in the direct care of patients in public hospitals and health centers in Addis Ababa were the study population.

### Selection criteria

All healthcare providers who were working in selected healthcare facility who have the qualification of physicians, health officers, nurses, midwives, and anesthesiologist personnel who work at least 6 months in the direct care of patients in public hospitals and health centers were included. Healthcare providers who were on annual and maternity leave during data collection period were excluded.

### Sample size determination and procedure

The sample size was determined using single population proportion formula, by considering the proportion of stethoscope disinfection after every use 50% (since there was no previous study in the study area). The following assumptions were used; 95% confidence interval (CI), 5% of marginal error, a design effect 1.5 and 10% for non-responder. Accordingly, a total of 576 healthcare providers were included.

A multi-stage sampling procedure was employed to select study participants. First, healthcare facilities were stratified by type into hospitals and health centers. Overall twenty one healthcare facilities were selected randomly using lottery method (3 hospitals and 18 health centers were selected from each stratum). Afterward, the calculated sample sizes were allocated proportional to size for each healthcare facility. Then, systematic random sampling was employed to identify the study population, after sampling frame preparation. And the first participant was selected randomly.

### Variables of the study and measurements

The dependent variable studied was stethoscope disinfection after every use (yes, no). Whereas, the independent variables includes; socio-demographic characteristics (age, sex, marital status, profession, educational level, and year of service); institutional related variables (training about infection prevention, working department, and availability of standard operating procedure (SOP) in working department) and individual related variables (awareness on Ethiopian infection prevention and patient safety (IPPS) guideline, attitude towards infection prevention, knowledge towards infection prevention and control of HAIs, self-reported infection prevention practice). Stethoscope disinfection practice after every use was measured using “yes” or “no” question. Accordingly, a score of “1” was assigned for acceptable disinfection practice (yes) and “0” for unacceptable (no) [18].

Knowledge about infection prevention and control of HAIs was measured using the cumulative score of 17 questions each with two possible response [i.e. “yes=1” or “no=0”]. A scoring system was used in which the respondent’s correct and incorrect answers provided for the questions were allocated “1” or “0” points, respectively. Knowledge scores were summed up to give a total knowledge score for each healthcare provider. The total score of knowledge questions ranging from 0 to 17 were classified into two categories of response: knowledgeable (if above the mean) and not knowledgeable (equal to or below the mean). Likewise, twelve questions were designed to assess participants practice regarding infection prevention. To analyze the practice, similar procedures were followed a score of 1 was assigned for each acceptable or “always practice response” and 0 for unacceptable, hence the total score of infection prevention practice ranged from 0 to 12. Accordingly, participant’s infection prevention practice was classified into two categories: safe practice (if above the mean) and unsafe practice (equal to or below the mean) [27].

There were twelve questions with Likert-type scale options ranging from “strongly agree’’ to ‘‘strongly disagree’’ to assess healthcare providers attitude towards infection prevention. Accordingly, mean value was used to classify infection prevention attitude as “favorable attitude towards infection prevention” if the score was equal or above the mean or “unfavorable attitude towards infection prevention” if the score was below the mean value [27]

### Data collection and quality control

A pre-tested interviewer administered questionnaire was used for data collection and four trained nurse were collect the data. The data collection tool was developed by reviewing related studies [18,15] and relevant literatures [20] and modified contextually. The data collection tool was first prepared in English and translated into Amharic (local language) then retranslated to English. Moreover, questionnaire was pre-tested on 5% of the actual sample size. The completeness, consistency, and accuracy of the collected data were examined on daily basis by two public health experts and by principal investigator.

### Data processing and analysis

After data collection, each questionnaire was checked for completeness, missing and edited for other errors. Statistical analyses were conducted with SPSS Statistics, version 20.0 (IBM, Armonk, NY, USA). A summary descriptive statistics were computed. Bivariate and multivariable logistic regression analyses were employed to identify factors associated with disinfection practice. Variables found significant at p-value 0.05 in bivariate analysis were included in to multivariable logistic regression analysis. Predicting power of variables in the final fitted model was checked by receiver observed characteristics (ROC) curve. The Hosmer and Lemeshow test was used for overall goodness of fit. Odds ratios with 95% confidence intervals (CI) were used to determine the strength of association between the outcome and explanatory variables. The statistical significance tests were declared at the p-value < 0.05 (S1 Table).

### Ethical statement

The study was ethically approved by Addis Ababa City Administration Health Bureau Institutional Ethical Review Board (IRB). Informed written consent was obtained from each healthcare provider after explaining the purpose of the study. The right of participants to anonymity and confidentiality was maintained.

## Results

### Socio-demographic characteristics

Five hundred forty six participants were take part in this study, for a response rate of 94.7%. Of these, 191 were male and 355 were female. Two hundred thirty seven, (43.4%) were in the age group between 26 and 30 years old. The mean (standard deviation [SD]) age of the respondents was 29.19 (SD ± 6.3) and majorities 60.6% of them were married. A higher proportion (61.9%) of the respondents was first degree and above and 68.1% of healthcare providers were nurses by profession (Table 1).

**Table 1:** Socio-demographic and other characteristics of the healthcare providers in healthcare facilities in Addis Ababa, Ethiopia 2016 (N=546).

| Variable | Characteristics | Number (n) | Percentage (%) |
| --- | --- | --- | --- |
| Age (year) | <25 | 164 | 30.0 |
|  | 25-30 | 237 | 43.4 |
|  | 31-35 | 75 | 13.7 |
|  | 36-40 | 28 | 5.1 |
|  | >40 | 42 | 7.7 |
| Sex | Male | 191 | 35.0 |
|  | Female | 355 | 65.0 |
| Marital status | Single | 331 | 60.6 |
|  | Married | 215 | 39.4 |
| **Profession** | Nurses | 372 | 68.1 |
|  | Health Officer | 65 | 11.9 |
|  | Midwives | 51 | 9.3 |
|  | Physicians* | 47 | 8.6 |
|  | Anesthesiologist | 11 | 2.0 |
| **Educational status** | First degree and above | 338 | 61.9 |
|  | Diploma | 208 | 38.1 |
| **Working department** | OPD, Emergency-OPD and Triage | 155 | 28.4 |
|  | Maternity, Delivery Gynecology and Obstetrics unit | 103 | 18.9 |
|  | Medical and Surgical Ward | 84 | 15.4 |
|  | Referral clinics | 42 | 7.7 |
|  | Pediatrics ward & NICU | 36 | 6.6 |
|  | Family planning & MCH | 55 | 10.1 |
|  | VCT, ART & TB-clinic | 42 | 7.6 |
|  | OR and Minor-OR | 29 | 5.3 |
| **Service year** | ≤ 5 | 432 | 79.1 |
|  | > 5 | 114 | 20.9 |
| **Infection prevention training** | Yes | 217 | 39.7 |
|  | No | 329 | 60.3 |
| **Availability of SOP** | Yes | 255 | 46.7 |
|  | No | 291 | 53.3 |

### Characteristics of institutional and individual conditions

In this study, high proportion 329 (60.3%) of the healthcare providers were trained on infection prevention. One hundred fifty five, (28.4%) of participates were working at OPD, Emergency-OPD and triage at the time of this survey. Two hundred and fifty five (46.7%) of healthcare providers reported that they had SOP targeted on infection prevention in there working department (Table 1).

### Stethoscope disinfection practice and other individual related characteristics

In this study, 217(39.7%) [95% confidence interval [CI]:35.9, 44.0%] of healthcare providers reported that they disinfect their stethoscope after every use. Whereas majority (60.3%)[95%CI: 56.2, 64.7%] of the respondent do not disinfect their stethoscope after every use, of these 189 (34.6%)[95%CI: 30.8,38.5%] of respondents never disinfect their stethoscopes and 84(15.4%)[95%CI:12.3,18.7%] and 56(10.3%)[95%CI: 7.7,12.8%] of healthcare providers disinfect their stethoscopes one a week or less often and one or two a day, respectively (Fig 1).

**Fig 1:**
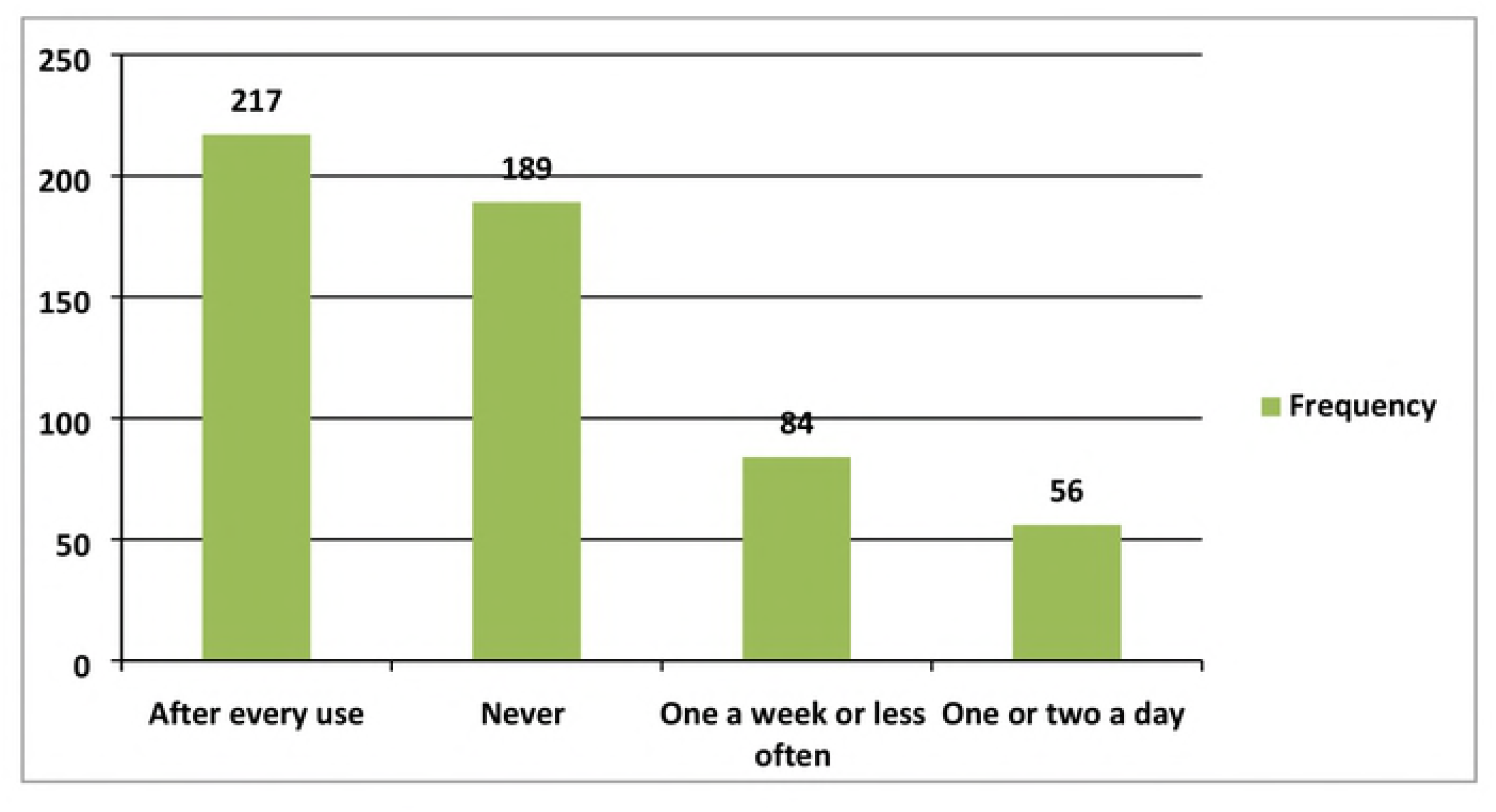
Frequencies of stethoscope disinfection among healthcare providers in Ethiopia, 2016 (N=546).

Four hundred and fifteen (76.0%) of healthcare providers belief transmission of infections occurs via stethoscopes. In addition, 308(56.4%) of healthcare providers were knowledgeable towards infection prevention and control of HAIs and 372(68.1%) have good infection prevention practice.

### Factors associated with stethoscope disinfection after every use

Table 2, 3 and 4 displays the results of bivariate and multivariate logistic regression analyses to identify factors of stethoscope disinfection after every use. In bivariate analyses; age, profession, working department, infection prevention training, awareness on infection prevention and patient safety (IPPS) guidelines, availability of standard operating procedures (SOP), belief that stethoscopes can transmit HAIs, attitude towards IP, knowledge on IP and control, and self-reported IP practice were significantly associated with stethoscope disinfection after every use. The final model was checked by Hosmer and Lemeshow test for the overall goodness of fit (0.714). In multivariate analyses, the odd of disinfection after every use was likely to be decreased by 79% among physician’s compared to nurses (Adjusted odds ratio [AOR]=0.21; 95% confidence interval [CI]: 0.09, 0.49). The odd of disinfection after every use was 1.93 times higher in healthcare providers who have awareness on infection prevention guideline than healthcare providers who did not have awareness (AOR=1.93; 95%CI: 1.31, 2.82). Among healthcare providers, the odds of disinfection after every use were significantly higher among who had favorable attitude towards infection prevention (AOR=1.73, 95%CI: 1.02, 2.93), and among those have safe infection prevention practice (AOR=3.79, 95%CI: 2.45-5.84) as compared with their counterparts.

**Table 2:** Socio-demographic factors bivarite analysis of factors associated with stethoscope disinfection after every use among healthcare providers in Ethiopia 2016.

| Variables | Stethoscope disinfection after every use |  | COR (95%CI) | P-value |
| --- | --- | --- | --- | --- |
|  | Yes (217) | No (329) |  |  |
| **Age** |  |  |  |  |
| <25 | 65 | 99 | 0.45(0.22-0.89)* | 0.02 |
| 25-30 | 87 | 150 | 0.39(0.20-0.77)* | 0.07 |
| 31-35 | 30 | 45 | 0.45(0.21-0.97)* | 0.04 |
| 36-40 | 10 | 18 | 0.38(0.14-1.02) | 0.05 |
| >40 | 25 | 17 | 1 |  |
| **Sex** |  |  |  |  |
| Male | 70 | 121 | 0.82(0.57-1.18) | 0.27 |
| Female | 147 | 208 | 1 |  |
| **Profession** |  |  |  |  |
| Nurses | 160 | 212 | 1 |  |
| Health Officer | 25 | 40 | 0.83(0.48-1.42) | 0.49 |
| Midwives | 19 | 32 | 0.79(0.43-1.44) | 0.43 |
| Physicians | 8 | 39 | 0.27(0.12-0.58)* | 0.00 |
| Anesthesiologist | 5 | 6 | 1.10(0.33-3.68) | 0.87 |
| **Educational level** |  |  |  |  |
| First degree and above | 135 | 203 | 1.02(0.72-1.45) | 0.90 |
| Diploma | 82 | 126 | 1 |  |
| **Year of service** |  |  |  |  |
| ≤ 5 | 164 | 268 | 0.70(0.46-1.07) | 0.09 |
| > 5 | 53 | 61 | 1 |  |

**Table 3:** Individual and institutional related factors associated with stethoscope disinfection after every use.

| Variables | Stethoscope disinfection after every use |  | COR (95%CI) | P-value |
| --- | --- | --- | --- | --- |
|  | Yes (217) | No (329) |  |  |
| **Working department** |  |  |  |  |
| OPD, Emergency-OPD and Triage | 57 | 98 | 1 |  |
| Referral clinics | 18 | 24 | 1.29(0.65-2.58) | 0.47 |
| Medical and Surgical Ward | 42 | 42 | 1.72(1.00-2.94)* | 0.04 |
| Pediatrics ward & NICU | 11 | 25 | 0.76(0.35-1.65) | 0.48 |
| Maternity, Delivery Gynecology and Obstetrics unit | 36 | 67 | 0.92(0.55-1.55) | 0.76 |
| OR and Minor-OR | 14 | 15 | 1.61(0.72-3.56) | 0.24 |
| FP, MCH, VCT, ART & TB-clinic | 39 | 58 | 1.16(0.68-1.94) | 0.58 |
| **Infection prevention training** |  |  |  |  |
| Yes | 101 | 116 | 1.59(1.13-2.26)* | 0.00 |
| No | 116 | 213 | 1 |  |
| **Do you have awareness on IPPS guidelines** |  |  |  |  |
| Yes | 150 | 184 | 1.76(1.23-2.53)* | 0.00 |
| No | 67 | 145 | 1 |  |
| **Availability of SOP** |  |  |  |  |
| Yes | 119 | 136 | 1.72(1.23-2.44)* | 0.00 |
| No | 98 | 193 | 1 |  |
| **Belief that stethoscopes can transmit HAIs** |  |  |  |  |
| Yes | 177 | 238 | 1.69(1.11-2.57)* | 0.01 |
| No | 40 | 91 | 1 |  |
| **Attitude towards IP** |  |  |  |  |
| Favorable | 192 | 262 | 1.96(1.19-3.22)* | 0.00 |
| Un-favorable | 25 | 67 | 1 |  |
| **Knowledge towards infection prevention and control of HAIs** |  |  |  |  |
| Knowledgeable | 136 | 172 | 1.53(1.08-2.18)* | 0.01 |
| Not knowledgeable | 81 | 157 | 1 |  |
| **Self-reported IP practice and control of HAIs** |  |  |  |  |
| Safe | 36 | 138 | 3.63(2.39-5.53)* | 0.00 |
| Unsafe | 181 | 191 | 1 |  |

SOP= standard operating procedure, IP= infection prevention, IPPS= infection prevention and patient safety

**Table 4:**
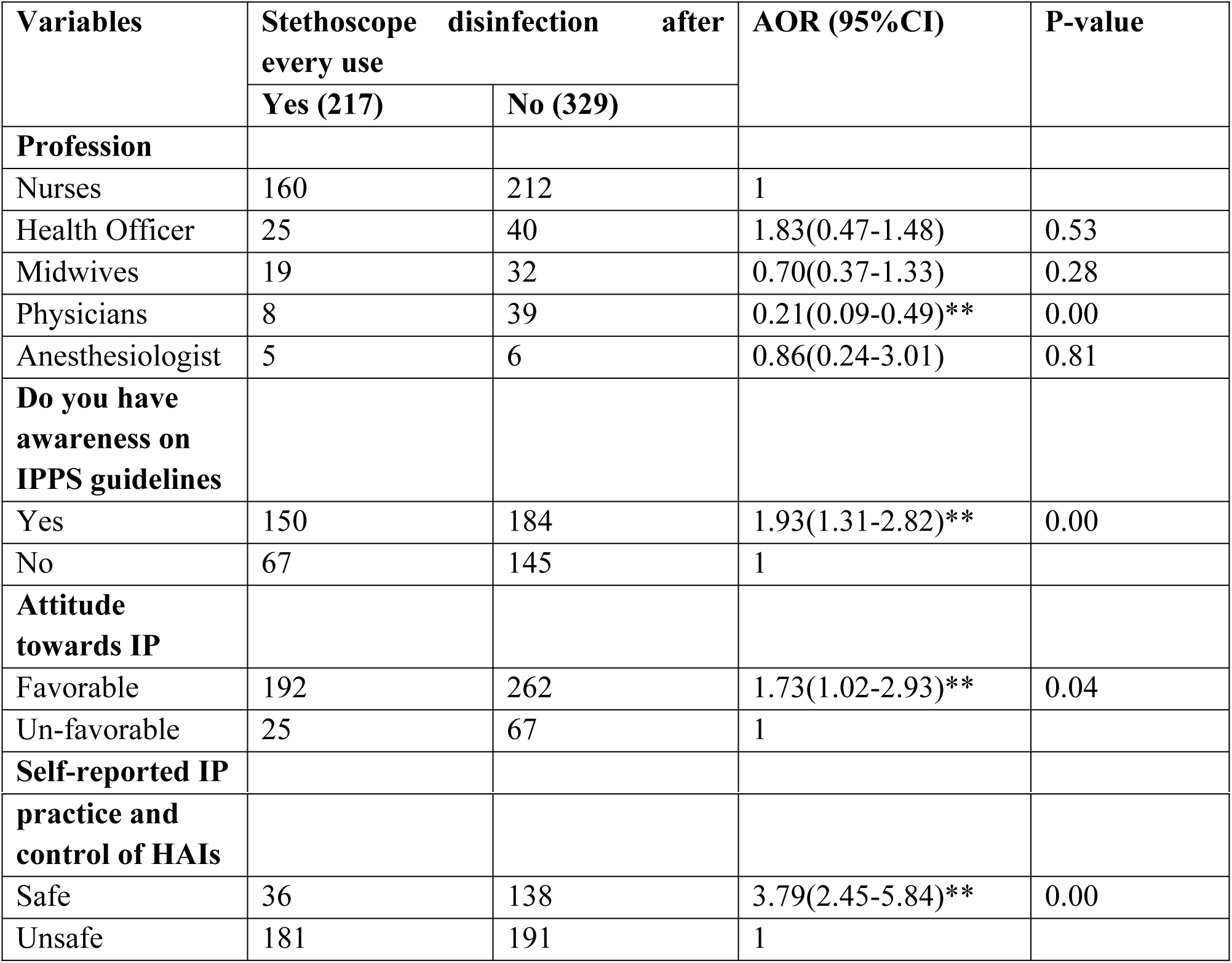
Multivariable logistic regression analysis of factors associated with stethoscope disinfection after every use α♣.

| Variables | Stethoscope disinfection after every use |  | AOR (95%CI) | P-value |
| --- | --- | --- | --- | --- |
|  | Yes (217) | No (329) |  |  |
| **Profession** |  |  |  |  |
| Nurses | 160 | 212 | 1 |  |
| Health Officer | 25 | 40 | 1.83(0.47-1.48) | 0.53 |
| Midwives | 19 | 32 | 0.70(0.37-1.33) | 0.28 |
| Physicians | 8 | 39 | 0.21(0.09-0.49)** | 0.00 |
| Anesthesiologist | 5 | 6 | 0.86(0.24-3.01) | 0.81 |
| **Do you have awareness on IPPS guidelines** |  |  |  |  |
| Yes | 150 | 184 | 1.93(1.31-2.82)** | 0.00 |
| No | 67 | 145 | 1 |  |
| **Attitude towards IP** |  |  |  |  |
| Favorable | 192 | 262 | 1.73(1.02-2.93)** | 0.04 |
| Un-favorable | 25 | 67 | 1 |  |
| **Self-reported IP** |  |  |  |  |
| **practice and control of HAIs** |  |  |  |  |
| Safe | 36 | 138 | 3.79(2.45-5.84)** | 0.00 |
| Unsafe | 181 | 191 | 1 |  |

## Discussion

Routine disinfection of stethoscopes after every use is one of the most important challenges among healthcare providers. In many developing countries, including Ethiopia healthcare provider’s stethoscopes disinfection practice scarcely addressed in scientific literatures. For this, the study assessed stethoscope disinfection practice and associated factors among healthcare providers.

In this study, more than three fifth (76.0%) of healthcare providers believe that stethoscope contamination can contribute to the transmission of infections. And, it was found that only a minority (39.7%) of healthcare providers reported that they disinfecting their stethoscopes after every use. The current finding was higher when compared to a study conducted in Mizan-Tepi University Teaching Hospital (southwest Ethiopia), in which only 6.45% of health professionals disinfect their stethoscope consistently [36]. Additionally, this finding also higher when compared to a study by Shiferaw et al among healthcare workers and medical student in Jimma University Specialized Hospital (Ethiopia) which reported, of the 176 stethoscopes studied, only 5 (2.8%) of the respondents (owners) reported they disinfect their stethoscope before and after examining each patient [34]. This finding could be attributed to different factors, including study setting, study participants, a difference in the definition of stethoscopes disinfection practice and other methodological concerns. Moreover, dissimilarity in awareness level of healthcare workers concerning infection prevention and stethoscopes disinfection could be another factor for this inconsistency. The present study also much higher than a study report by Dabsu et al [35], in Tikur Anbessa Specialized Referral Hospital (Ethiopia), which reported 8.5% of the study healthcare workers clean their stethoscopes between patient examinations.

The current finding indicated a significant number of healthcare providers rarely disinfect their stethoscope. In connection, different studies from different parts of the globe reported infrequent disinfection practice habits, a study from Nepal only (6.89%) of healthcare workers disinfecting after every use [14], in USA 24% of healthcare workers only disinfect their stethoscopes after every patient [18], in Turkey cleaning of stethoscopes with various disinfectants at certain intervals was reported 50.4% [38], in Pakistan, a considerably lower prevalence of stethoscope cleaning was observed 37.7% [39], and in Nigeria 87.9% of the participants did not clean their stethoscopes after examining each patient [40].

However, the current finding was lower than the finding from Serbia, in which (80.8%) of the respondent clean their stethoscopes [41], and a study from French, reported (82%) of study participants clean their stethoscopes regularly or from time to time [42]. Additionally, the current finding was much lower than a study from Scotland, in which 78.6% of respondents clean their stethoscopes [43]. These differences might be attributed to a distinctive background of study participates, medical practice, and awareness level dissimilarity. A choice of study population could be another possible explanation for this differences, as in the Serbia survey respondents were four and sixth year medical students and in the cases of French and Scotland survey respondents where medical students.

In connection, different literatures extensively reported contamination of stethoscope is likely, if stethoscope not disinfected regularly. A literature search by O’Flaherty et al across several databases for relevant studies on stethoscopes showed that stethoscopes were consistently harbor bacteria and the mean rate of stethoscope contamination across 28 studies was 85.1% (range: 47‒100%) [44]. A prospective study conducted in Swiss university teaching hospital to compare the contamination level of physicians’ hands and stethoscopes provide a strong evidence of the potential for stethoscope-mediated transmission of microorganisms and the need to systematically disinfect stethoscopes after each use [45].

A study by Pal et al [46], try to identify factors associated with stethoscope disinfection and reported apprehension of damaging stethoscopes and lack of knowledge regarding good disinfectant were the underlying causes that prevent cleaning of the stethoscopes. In the same way, this study identified factors associated with stethoscope disinfection. Healthcare providers who have awareness on infection prevention guideline were more likely to disinfect their stethoscope than their counterpart healthcare providers. This could be due to the fact that as the healthcare provider’s exposure to such guidelines increase, healthcare providers are frequently exposed to appropriate stethoscopes disinfection practices and become more compliant. According to the Healthcare Infection Control Practices Advisory Committee Guideline for Disinfection and Sterilization in Healthcare Facilities recommended appropriate cleaning of stethoscopes with 70% ethyl or isopropyl alcohol after every use is recommended [20]. In addition with this guideline, the Ethiopian infection prevention and patient safety guideline also recommended disinfection of stethoscopes after every use [47].

It was found that healthcare providers who had favorable attitude towards infection prevention were 1.73 times more likely to disinfect their stethoscope after very use than those healthcare providers who had unfavorable attitude. The current finding was in agreement with a study conducted by Gazibara et al [41], which reported a positive correlation between a higher frequency of stethoscope cleaning and the stronger positive notion that a stethoscope should be cleaned. In line with this finding a study conducted in Addis Ababa also reported a significant positive association between positive attitude towards infection prevention and good infection prevention practices [27].

In addition, this study reveled that infection prevention practice is another factor significantly associated with the practice of stethoscope disinfection after every use. Healthcare providers who had safe infection prevention practice were four times more likely to disinfect their stethoscope as compared to those who had unsafe infection prevention practice. This can be explained by the fact that, disinfection of medical equipments that comes into contact with patients is one of the core principle of infection prevention and those healthcare workers who had good compliance towards infection prevention may have better awareness and compliance towards stethoscope disinfection.

Furthermore, different studies try to identify factors associated stethoscope disinfection. A study by Wood et al [48], identified concern for damage of stethoscope, lack of time and lack of knowledge regarding best cleaner were identified as barrier of stethoscope cleaning [48]. A study by Hyder also suggests that promotion of stethoscope cleaning may have a positive effect on cleaning compliance in hospital wards. In addition, it was showed that a history of receiving information on stethoscope cleaning has been one of the strongest predictors of stethoscope hygiene [39].

In the present study, physicians were less likely to disinfect their stethoscope after every use compared to nurses. This finding was supported by different similar studies. A study by Shiferaw et al [34], in their study reported all licensed doctors (specialist, resident and general practitioner) didn’t disinfect their stethoscope regularly. Consequently, 98% of the studied stethoscope diaphragms were contaminated. Moreover, other studies documented doctors disinfect their stethoscope irregularly [48,49]. The current finding was also in consistent with other findings, where nurses reported to have good disinfection practice than doctors [49, 50]. Other previous conducted studies also indicated that about 97 to 100% of doctors did not follow a standard disinfection protocols [34,49,51]. In light of this finding, a study by Uneke et al described stethoscopes used by physicians were more contaminated than those used by other health workers [40].

### Limitations of the study

This study has several limitations. Due to the cross-sectional nature of this study design temporal relationships cannot be established. The study did not perform any direct observation of disinfection practices to externally validate healthcare provider’s responses. Therefore, social desirability bias is likely. The other limitation of this study is that the generalization of findings limited to public healthcare facilities. It was conducted in public healthcare facilities, thus limiting its generalizability to such settings. One additional limitation of the current study was it did not collect samples from the diaphragm, other parts of the stethoscope, such as plastic ear pieces to identify bacterial contamination. Lastly, subsequent observational studies are required to determine actual practice and to investigate if cleaning of stethoscopes leads to a reduction in HAIs in healthcare settings.

## Conclusions

The present study confirmed that only a small proportion of healthcare providers disinfect their stethoscopes after every use. Consequently, the possibility of stethoscopes contamination with pathogenic microorganisms is likely, and contributes to elevate the risk of contracting HAIs. Factors such as awareness on infection prevention guidelines, favorable attitude towards infection prevention and safe infection prevention practice and control of HAIs were the independent predictors of stethoscopes disinfection after every use. The study also reveled that physicians were less likely to disinfect their stethoscope compared to nurses. Hence, implementation of effective training on stethoscope disinfection along with increasing awareness of healthcare providers on infection prevention practice may improve stethoscope disinfection after every use. In addition, access to infection prevention guidelines in all healthcare settings should be strengthening to enhanced adherence towards stethoscope disinfection.

## Acknowledgements

I would like to thank all participants involved in the undertakings of the study and Addis Ababa health city administration health Bureau.

## Supporting information

S1 Table: Description of socio-demographic, institutional, and individual related variables included in this analysis.

